# Catalytic activity of KMT5B promotes cilia tuft formation without affecting chromatin accessibility of ciliary genes

**DOI:** 10.1101/2025.03.18.644017

**Authors:** Janet Tait, Carmen Marthen, Barbara Hoelscher, Tobias Straub, Ralph Rupp

## Abstract

Multiciliated cells are a specialized cell type found in the brain, reproductive tract and respiratory tract of mammals. Multiciliated cells resembling those of the mammalian lung can also be found on the surface of the epidermis of tadpole stage Xenopus embryos, making the frog an ideal model organism to study this cell type. KMT5B/suv4-20h1 is a histone methyltransferase that writes the dimethyl mark on histone 4, lysine 20 (H4K20). We have previously shown that the multiciliated cells of embryos lacking KMT5B have depleted cilia and a reduced actin cap, and that knockdown of both KMT5B and KMT5C/suv4-20h2 leads to downregulation of ciliogenic genes. Here we further tease out the independent function of KMT5B in multiciliogenesis, and show that single knockdown of KMT5B, but not KMT5C leads to aberrant transcription and the downregulation of cilia genes. This phenotype is ameliorated by overexpression of catalytically active PHF8, an H4K20me1 demethylase, while a hormone inducible variant of multicilin (MCI), the master regulator of cilia tuft formation, has no effect. Notably, the expression of key transcription factors of ciliogenesis is unaffected by KMT5B depletion, and the transcriptional effect of KMT5B depletion dominates the response to ectopic multicilin. Finally, ATAC seq analysis in animal caps reveals that knockdown of KMT5B results globally in very few peaks with differential activity and does not compact chromatin at ciliary genes. Taken together, this suggests that KMT5B regulates multiciliated cells through an alternative pathway to the canonical MCI-driven multiciliogenic program.

## INTRODUCTION

Cellular differentiation is a dynamic process by which cells progress from pluripotency into their specific cell fate. Sometimes they acquire specialized structures and employ complex gene regulatory networks in the process of differentiation. Multiciliated cells are a highly specialized and post-mitotic cell type found in the mammalian brain, fallopian tubes, and bronchi (Bustamante-Marin and Ostrowski). These cells contain hundreds of motile cilia that beat in a coordinated manner to generate fluid flow. Multiciliated cells resembling those of the mammalian lungs are also found on the surface of Xenopus embryonic epidermis (Walentek).

Multiciliated cells are specified by Notch signaling, which selects for MCC identity by inducing the expression of the geminin proteins multicilin (MCI), the so called “master regulator” of ciliogenesis, and gemc1 in humans (Meunier and Azimzadeh; Ma et al.; J. L. Stubbs et al.). From there, a cascade of transcription factors tightly controls ciliogenesis. Multicilin binds E2F4/5 transcription factors, which induces cell cycle exit in favour of differentiation. C-MYB, an S-phase protein, and CCNO, a cyclin-like protein, coordinate the formation of basal bodies, the specialized centrioles that nucleate cilia (Ma et al.; J. L. Stubbs et al.). Simultaneously, RFX family transcription factors alongside FOXJ1 control cilia formation and the formation of the actin cap, a lattice of actin meshwork at the apical surface of multiciliated cells (Quigley and Kintner). Our lab previously demonstrated an unexpected link between Histone 4 Lysine 20 (H4K20) methylating enzymes and multiciliated cells in the Xenopus embryonic epidermis (Angerilli et al.).

Histone methyltransferases are responsible for writing methyl marks on specific amino acid residues on histone tails. H4K20 can be unmethylated, mono-, di-, or trimethylated. The marks are present at different abundancies on the chromatin, with approximately 10% unmethylated, 10% H4K20me1, 80% H4K20me2 – the highest abundance of any histone modification on the chromatin, and less than 1% H4K20me3 (Corvalan and Coller; Pokrovsky et al.; Yang et al.; Young et al.). This mark is written in a cell cycle-dependent manner and there are a number of enzymes that write at this site. H4K20me1 is written by SET8/PR-SET7 in G2-phase and M-phase, before proteolytic degradation of SET8 in late G1 phase (Oda et al.). H4K20me1 is then largely converted into H4K20me2 by KMT5B/SUV4-20H1 and KMT5C/SUV4-20H2 and further on to H4K20me3 by KMT5C (Eid et al.; Schotta et al.). H4K20 demethylases have also been identified. PHF8 converts H4K20me1 into unmethylated H4K20 (Liu et al.; Qi et al.), while RSBN1 has been shown to demethylate H4K20me2 (Brejc et al.), and recently hHR23b has been shown to have activity towards all H4K20me states in vitro (Cao et al.).

H4K20 methylating and demethylating enzymes also play critical roles in controlling gene expression during embryonic development. SET8 depletion is embryonic lethal in mice and *Drosophila* (Schotta et al.; Oda et al.). Despite high sequence homology between KMT5B and KMT5C, they have diverse functions and roles in development. KMT5B is ubiquitously expressed during mouse development, while KMT5C abundance is lower and more tissue-specific. KMT5B knockout also results in perinatal lethality, while KMT5C knockout mice develop normally (Schotta et al.).. In *Xenopus*, double knockdown of KMT5B and KMT5C results in ectodermal defects including craniofacial abnormalities, loss of melanocytes and reduced eye structures (Nicetto et al.; Angerilli et al.). KMT5B is also a well-known autism spectrum disorder risk factor gene (Wickramasekara and Stessman). PHF8 knockdown also has developmental consequences. In zebrafish, PHF8 knockdown leads to craniofacial defects as well as neurodevelopmental defects in the neural tube, and PHF8 is linked to Siderius X-linked intellectual disability (Qi et al.).

We previously showed that double knockdown of KMT5B and KMT5C have strong phenotypic and transcriptional effects on *Xenopus* multiciliated cells, and further, that knockdown of KMT5B alone can recapitulate the phenotypic changes in multiciliated cells (Angerilli et al.). Knockdown MCCs have much fewer, shorter cilia, a reduced apical actin cap, and in some cases, clumped basal bodies. Here we investigate this ciliogenic phenotype further and show that KMT5B knockdown alone has strong effects on gene expression, particularly expression of ciliogenic genes, in contrast to KMT5C, and that KMT5B knockdown drives transcriptional changes even in presence of exogenous multicilin, using multicilin overexpressing animal caps (ectodermal organoids). Additionally, we show that rescue of cilia by PHF8 relies on catalytic activity of the enzyme. Lastly, we profile the effect of KMT5B knockdown on chromatin accessibility. Taken together, our data sheds further light on the role of KMT5B in multiciliated cells.

## RESULTS

### Knockdown of KMT5B results in transcriptional changes

We performed confocal microscopy on embryos in which either KMT5B, or KMT5B were knocked down by translation-blocking morpholinos (Figure 1A). Confirming previous wholemount immunocytochemistry results, we see that knockdown of KMT5C has no effect on cilia or the actin cap, while knockdown of KMT5B results in fewer, shorter cilia, and a depleted actin cap as described for KMT5B/C double knockout (Angerilli et al.). To determine the individual effects of depleting KMT5B and KMT5C on gene expression, we compared the gene expression profiles of KMT5B and KMT5C knockdown animal cap organioids. Animal caps are explants taken from the blastocoel roof of blastula stage embryos and they differentiate by default into a mucociliary epithelium. Using animal caps instead of whole embryos results in a simplified tissue in which only 5 cell types are present instead of the hundreds of cell types of a whole embryo. Our first observation was that very few genes were misregulated by KMT5C knockdown. While more than 2000 genes are misregulated in KMT5B knockdown animal caps, only 83 genes were misregulated in KMT5C knockdown animal caps (Figure 1B, C). We attempted to perform GO analysis on the misregulated genes upon KMT5C knockdown and found no enriched GO categories for the upregulated genes and only three enriched GO terms in the downregulated genes: “regulation of transcription by RNA polymerase II”, “circadian rhythm”, and “circadian regulation of gene expression”, each category containing very few genes. This is likely due to the overall small cohort of misregulated genes. That KMT5C has such a weak effect on transcription is surprising because we previously showed that both KMT5B and KMT5C are responsible for writing H4K20me2 in the *Xenopus* embryo, and KMT5C further writes H4K20me3, a key heterochromatic mark.

**Figure 1:**
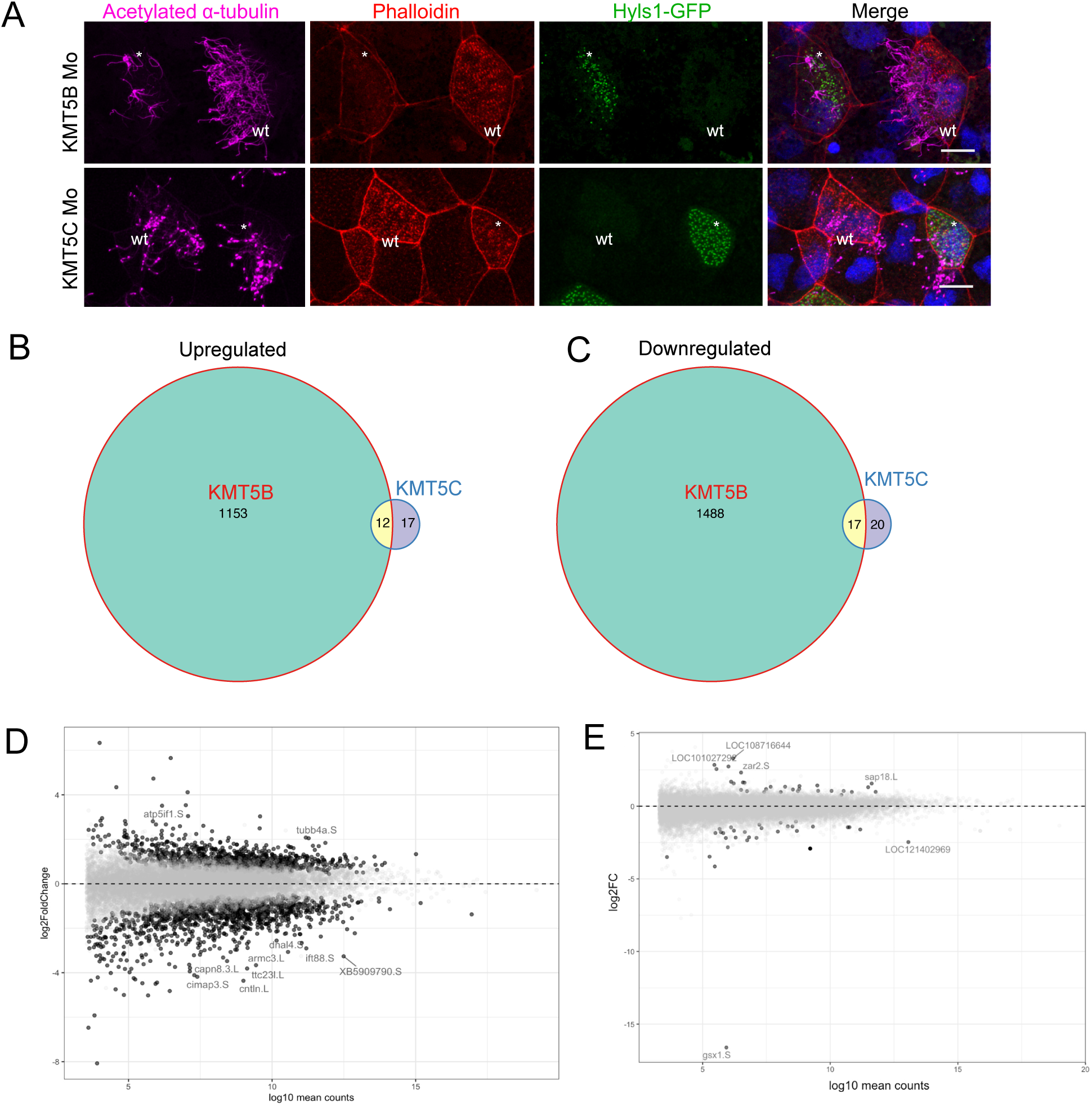
Knockdown of KMT5B, but not KMT5C leads to phenotypic and transcriptional changes. A) 4 channel confocal images depicting single knockdowns of KMT5B or KMT5C in *Xenopus laevis* embryos. Cilia are magenta (acetylated α-tubulin), basal bodies are green (hyls1-GFP), actin meshwork and cell borders are red (phalloidin), and DNA is blue (DAPI). Embryos are injected in one of two ventroanimal blastomeres at the 8-cell stage, giving rise to mosaic embryos in which uninjected (labelled wt in white) and knockdown (white asterisks) multiciliated cells can be visualized in the same field of view. B, C) Venn diagrams showing the number of genes that are upregulated (B) or downregulated (C) upon knockdown of KMT5B or KMT5C in *Xenopus laevis* animal caps by RNA-seq. D,E) MA plot showing log2fold change (x-axis) and log10 mean counts (y-axis) in KMT5B (D) or KMT5C (E) knockdown animal caps by RNA-seq.

KMT5B misregulation, on the other hand led to 1165 upregulated genes and 1505 downregulated genes (Figure 1B, C). Immediately, a ciliogenic connotation stuck out in the misregulated genes. For example, among the top misregulated genes were cntln, a centriolar linked protein that binds microtubules (Jing et al 2016), intraflagellar transport protein IFT88, and dnal4, a dynein protein found in motile cilia (Figure 1D). To our surprise, we found a tubulin protein, tubb4a, to be one of the most upregulated genes. However, tubb4a is one of two ß-tubulins that contain a C-terminal motif that allows them to interact with cilia (Figure 1E). However, while knockout of tubb4b leads to severe ciliogenic defects in the MCCS of the mouse airway, tubb4a depletion does not affect cilia tuft formation. Tubb4b is also downregulated upon KMT5B depletion. Tubb4a may also play a role in promoting proliferation (zhang et al, 2023).

We went on to perform GO analysis, and confirmed a strong ciliogenic connotation in the downregulated genes. The top misregulated GO terms include “cilium movement”, “cilium organization”, and “cell projection” (Figure 2A). Some of the upregulated GO terms related to mRNA processing and macromolecule localization. However, we were also interested to see that some terms related to mitosis and the cell cycle could be found in the upregulated GO terms, including “mitotic cell cycle” and “microtubule cell cytoskeleton organization involved in mitosis” (Figure 2B). Overall, this seems to suggest that KMT5B knockdown favours proliferation over formation of cilia.

**Figure 2:**
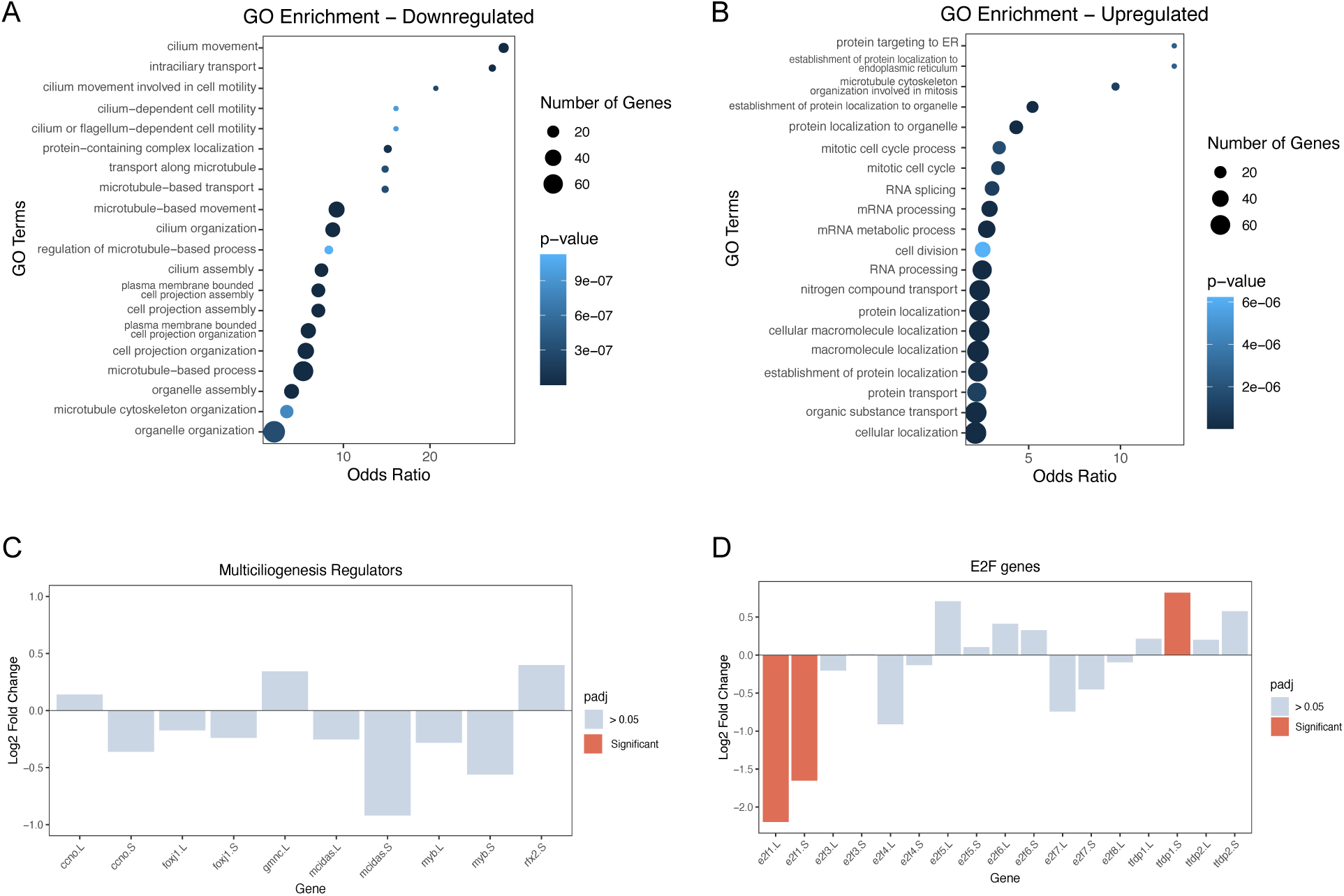
KMT5B regulates ciliogenesis through an alternative pathway. A,B) GO analysis of genes that are downregulated (A) or upregulated (B) upon KMT5B knockdown. Upregulated genes largely relate to mRNA processing and macromolecule localization, while downregulated genes relate to cilia and microtubules. C) Log2 fold change of multiciliogenic regulators upon KMT5B knockdown. None of the major known ciliogenic regulators are significantly misregulated based on RNA-seq results. D) Log2 fold change of E2F genes upon KMT5B knockdown. Only the L and S copies of cell cycle-related e2f1 is downregulated, while tfdp1.S is upregulated.

### Ciliogenic regulators are unaffected by KMT5B knockdown

What could explain this striking downregulation of ciliogenic genes? We reasoned that knockdown of KMT5B could lead to the downregulation of a key ciliogenic transcription factor, subsequently misregulating the rest of the ciliogenic differentiation pathway. Of particular interest are members of the E2F transcription factor family, some of which are known to play a role in multiciliogenesis. E2F transcription factors have a variety of distinct functions despite the fact that they share high sequence homology. They can be split into cell cycle-activating and cell cycle-repressing E2Fs. The cell cycle-activating E2Fs are E2F1, -2, and -3, while the cell cycle repressing E2Fs are E2F4-8. Multicilin, the master regulator of ciliogenesis, directly binds E2F4/5 and this binding is required for basal body biogenesis (Quigley and Kintner). Recently, E2F7 has been identified to play a key role in multiciliogenesis by attenuating DNA replication levels to favour MCC differentiation over proliferation (Choksi et al.).

We looked at the change of expression levels of E2F transcription factors and their binding partners (Figure 2D). Both the .L and .S homeolog of E2F1 were significantly misregulated. E2F1 is primarily involved in cell cycle regulation and has not been shown to play roles in multiciliogenesis. Additionally, tfdp1, binding partner of the cell cycle activating E2Fs, was significantly upregulated, with a log2 fold change of 0.82. The mRNA levels of other ciliogenic transcription factors 2 2like foxj1, rfx2 or myb were not significantly changed (Figure 2D). Taken together, the fact that neither the regulators of ciliogenesis, nor the majority of E2F transcription factors are misregulated suggests that KMT5B is operating through a different gene expression pathway.

### Catalytic activity of PHF8 rescues ciliogenic phenotype

We previously showed that ciliogenic defects can be partially rescued by PHF8 overexpression, but wanted to determine whether this rescue depends on catalytic activity. To test this, we obtained two hyperactive, truncated human PHF8 clones including amino acids 1-489 (Fortschegger et al.). We injected either wt cDNA (489 wt) or catalytically inactive (489 c.i.) phf8 mRNA into one of two blastomeres of a two cell-stage embryo alongside LacZ mRNA to act as a lineage tracer. We fixed the embryos at tadpole stage and performed whole mount immunocytochemistry against acetylated α-tubulin and performed β-galactosidase staining, which stains LacZ positive regions blue. We also injected luciferase mRNA to act as a non-specific mRNA control. In each condition the mRNA was co-injected with either control morpholino or KMT5B morpholino (Figure 3A). In total, 95.6% of KMT5B MOs and 86.6% of KMT5B MOs + 489 c.i. PHF8 embryos showed cilia tuft defects, while only 36% of KMT5B MOs + 489 wt PHF8 embyos showed cilia tuft defects. This indicates that the ciliogenic rescue depends on enzymatic activity of PHF8.

**Figure 3:**
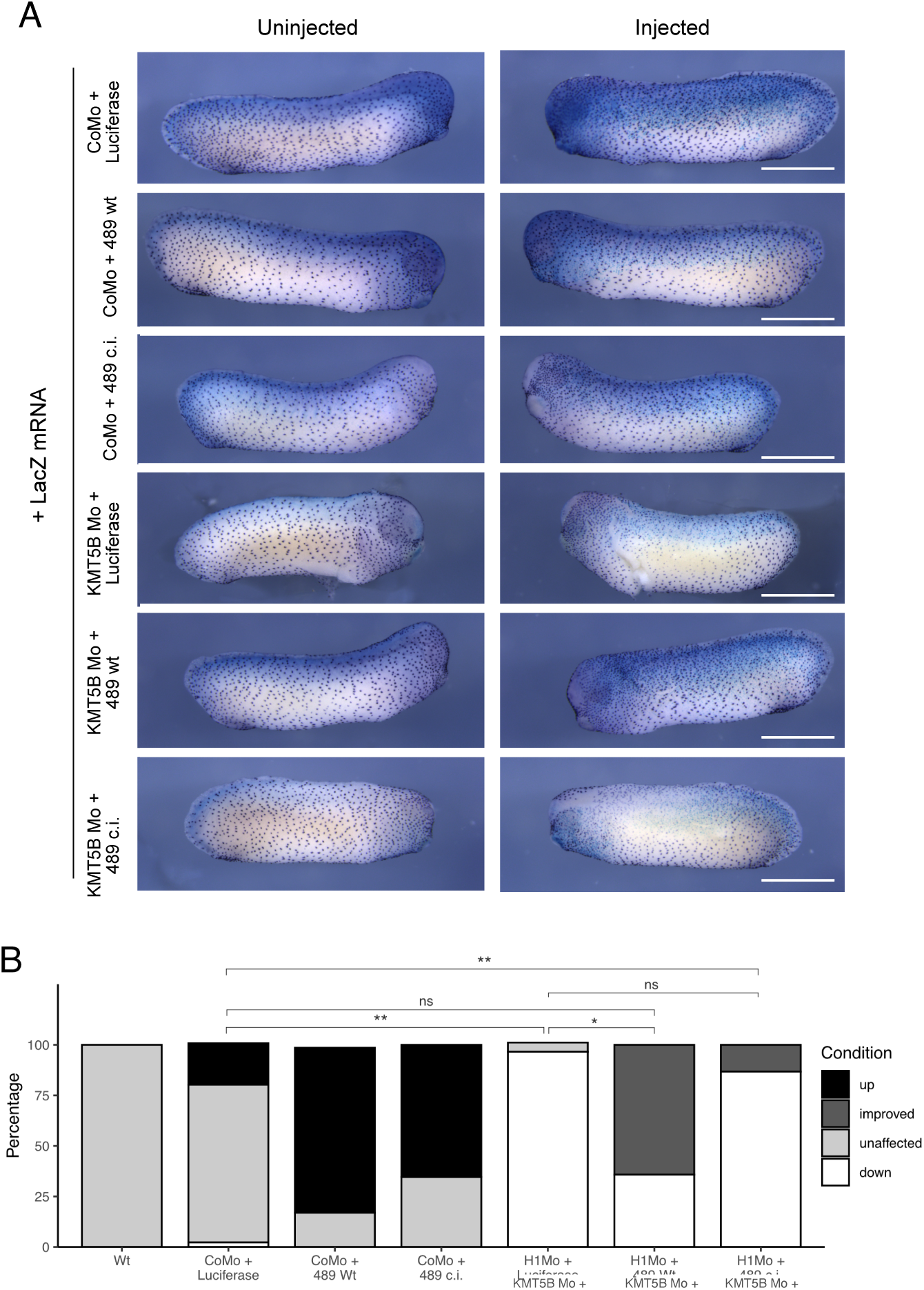
Catalytic activity of PHF8 rescues the ciliogenic phenotype. A) Representative immunocytochemistry images from tailbud stage embryos injected with KMT5B Mo or CoMo and mRNAs luciferase or human PHF8 variants (489 wt/489 c.i.). Scale bars = 1 mm (whole embryo), or 200 μm (inserts) and n = 3 biological replicates. B) Quantification of A). Percentage of embryos either affected by morpholino injections (H1Mo) or rescued by PHF8 mRNA injection. Wt PHF8 mRNA (489 Wt) rescues significantly, while catalytically inactive PHF8 mRNA (489 c.i.) shows no significant difference when compared to H1Mo. Significance is indicated by asterisks; * = p < 0.05, ** = p < 0.01, ns = not significant. N=4 biological replicates.

Additionally, we noted that both truncated clones affected the spacing of MCCs in wildtype embryos. Cilia tuft density increased in 81% of 489 wt embryos and 65% of 489 c.i. embryos co-injected with CoMo (Figure 3B). This result, which is independent of PHF8 enzymatic activity, suggests an involvement of PHF8 in MCC specification, rather than MCC differentiation. Further experiments are needed to corroborate this observation.

### Multicilin overexpression results in more MCCs, but does not rescue KMT5B phenotype

We wanted to further establish that KMT5B operates outside of the multicilin-driven gene expression pathway, so we decided to deplete KMT5B in the context of multicilin overexpression. Multicilin overexpression has also been shown to induce other epidermal cells to adopt an MCC-like fate (Quigley and Kintner). In a typical *Xenopus* epidermis, MCCs are one of 5 cell types, representing approximately 20% of cells (Plouhinec et al.). Multicilin overexpression has the added benefit of driving more cells to become multiciliated, giving a more representative profile of MCCs. We used a hormone-inducible multicilin construct containing the ligand binding domain of the human glucocorticoid receptor (MCI-hGR) that has been previously validated (Quigley and Kintner). To test the effect of KMT5B in multicilin over-expressing embryos, we injected 60 pg of MCI-hGR into one ventroanimal blastomere of a 8-cell stage embryos. We co-inject hyls1-GFP to act as a lineage tracer, indicating which MCCs are wt and which have been affected by multicilin overexpression with or without KMT5B knockdown. At early gastrula stage (NF11), we induced the embryos using 10 µl dexamethasone. Then, at tailbud stage we fixed the embryos and performed whole mount immunocytochemistry against acetylated alpha-tubulin (For a schematic of the method, see Figure S1A). We see that in embryos injected with control morpholino + MCI-hGR, almost the entire surface of the embryo is covered with multiciliated cells. By contrast, in KMT5B morpholino + MCI-hGR injected cells appear to adopt a multiciliogenic fate – evident by the multiplied GFP-tagged basal bodies, but they are not able to form cilia (Figure 4A). Due to this finding, we hypothesize that this is truly a phenotype relating to cilia formation, rather than a cell-specification phenotype. These cells can still undergo early stages of multiciliogenesis like amplification of centrioles and emergence into the surface of the embryo, but they cannot form cilia.

**Figure 4:**
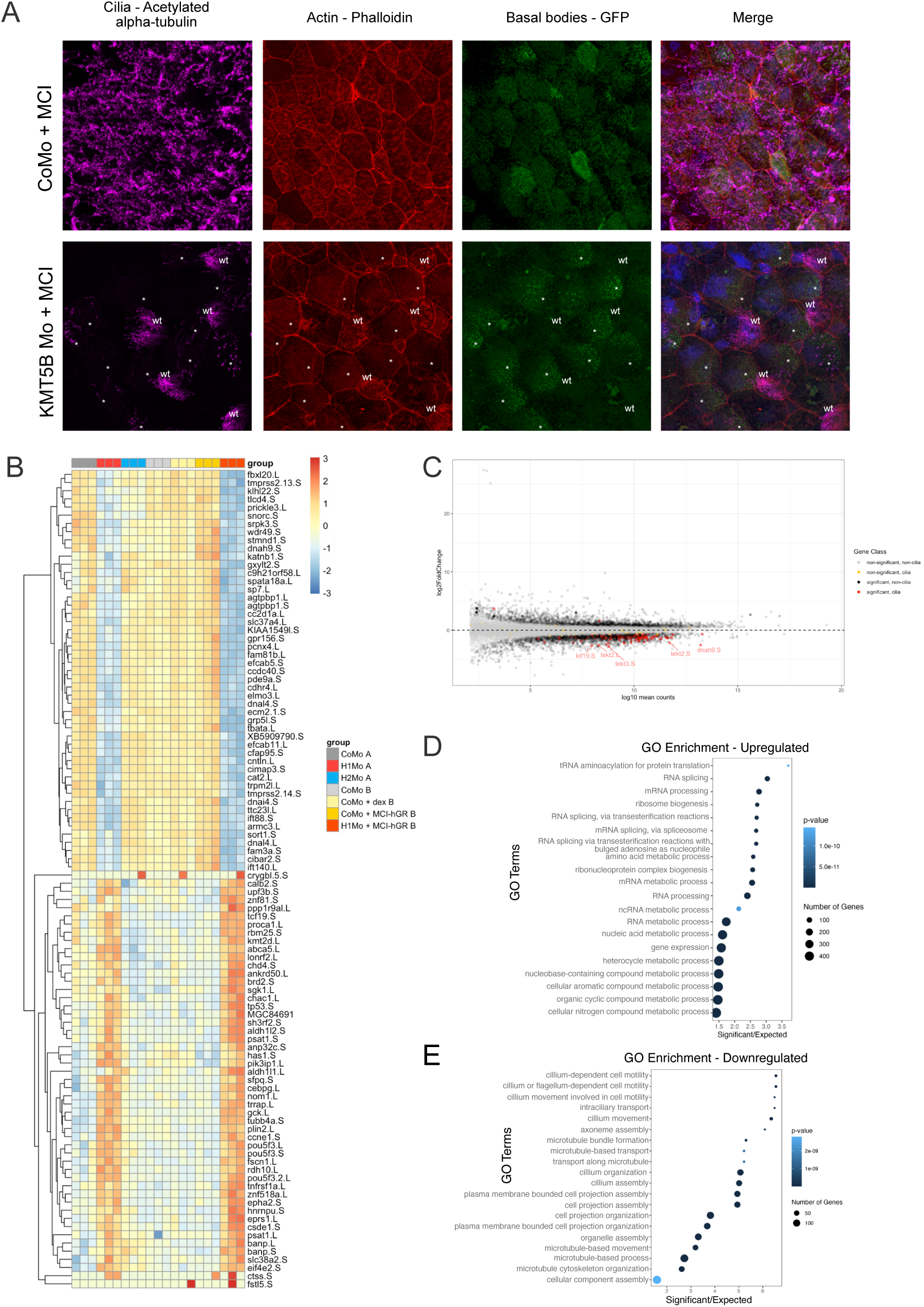
KMT5B knockdown drives transcriptional and phenotypic changes even in the presence of MCI-hGR. A) 4-channel confocal microscopy images showing Xenopus laevis embryos injected with control morpholino (CoMo) + MCI-hGR or KMT5B morpholino + MCI-hGR. Cilia are magenta (acetylated α-tubulin), basal bodies are green (hyls-GFP), actin meshwork and cell borders are red (phalloidin), and DNA is blue (DAPI). Embryos are injected in one of two ventroanimal blastomeres at the 8-cell stage, giving rise to mosaic embryos in which wt and knockdown multiciliated cells can be visualized in the same field of view. Uninjected cells are labelled in white as “wt”, while injected cells are indicated with an asterisk (*). B) Cluster analysis heatmap of top responding genes across all conditions and replicates. Genes group into two main clusters: genes downregulated or upregulated upon suv4-20h1 knockdown (in the absence or presence of MCI-hGR). Group A includes replicates from the single knockdown RNA seq experiment, and group B includes replicates from the MCI-hGR containing RNA seq experiment, and samples are batch corrected between experiments. C) MA plot showing all genes (light grey), significantly misregulated genes (dark grey), cilia genes (yellow), and significantly misregulated genes (red). D, E) GO analysis showing the upregulated (D) and downregulated (E) gene categories upon MCI-hGR overexpression. The downregulated gene categories are largely related to ciliogenesis and microtubule assembly.

We wanted to understand what transcriptional changes were underlying this phenotype, so we performed RNA-seq in animal caps. We injected embryos at the 2-cell stage with MCI-hGR and the morpholino of interest. At blastula stage (NF9) we dissected animal caps, which we then induced with dexamethasone at gastrula stage (NF11). We harvested the embryos at neurula stage (NF16) and performed RNA-seq on pools of 10 animal caps. Alongside this, we performed quality control ICCs in embryos to make sure that both the KMT5B knockdown phenotype and the multicilin overexpression phenotype were present in the batches of animal caps selected for sequencing. We also staged animal caps based on these sibling embryos.

Overexpression of MCI with control morpholino injection (CoMo + MCI-hGR) led to the misregulation of 591 genes (Figure S1B, C). Of this, 525 genes were upregulated and only 66 genes were downregulated. In line with our expectations, GO analysis of the upregulated gene categories revealed that categories related to MCCs were upregulated including “centriole assembly”, “cilium assembly”, and “cilium organization” (Figure S1D). On the other hand the downregulated genes were enriched for GO categories related to the cell cycle including “DNA replication”, and “DNA replication initiation”, as well as a category related to a different cell lineage, “mesoderm development”, and “negative regulation of cell differentiation” (Figure S1E). The downregulation of genes related to negative regulation of cell differentiation is unsurprising because MCCs are post-mitotic and terminally differentiated, and so DNA replication and its related processes should be restricted.

### Transcriptional changes are driven by KMT5B knockdown

MCI-hGR overexpression results in upregulation of ciliogenic genes, while KMT5B knockdown results in downregulation of ciliogenic genes. We wondered whether concurrent MCI-hGR overexpression and KMT5B depletion would have competing effects on ciliogenic gene expression. To better understand how these regulators affect the ciliogenic gene expression program, we performed a combined analysis of the two RNA-seq experiments, Group A (independent knockdowns of KMT5B and KMT5C) and Group B (MCI-hGR overexpression experiment). We found two major clusters, both defined by the behaviour of genes in response to KMT5B knockdown (Figure 4B). One cluster that is defined by genes that are downregulated in KMT5B knockdown and KMT5B knockdown + MCI-hGR, and a second cluster that is defined by genes that are upregulated in KMT5B knockdown and KMT5B knockdown + MCI-hGR. This indicates that KMT5B knockdown, and not MCI-hGR overexpression is the driver of these transcriptional changes.

We also performed GO analysis on KMT5B knockdown + MCI-hGR overexpressing animal caps. In the upregulated genes, we see categories relating to RNA processing and a number of metabolic processes, however unlike in the KMT5B knockdown, there is no cell cycle connotation. The downregulated genes are once again dominated by ciliogenic GO terms. This is despite the fact that ciliogenic genes are upregulated in MCI-hGR overexpressing animal caps, and the fact that MCI-hGR overexpression results in a higher density of multiciliated cells. Taken together, this suggests that regulation of multiciliogenesis by KMT5B proceeds either through an alternative pathway to, or downstream of the MCI-initiated transcription factor cascade.

### Effect of KMT5B on chromatin accessibility

H4K20me1 has been shown to have roles in chromatin compaction through its reader protein L3MBTL1, and through interactions with the C-terminus of H2A on neighboring histone tails. Could the downregulation of ciliogenic genes result from a loss of chromatin accessibility at target genes? We decided to use ATAC-seq to address this question. ATAC-seq peaks indicate open regions of the chromatin, and novel or changing peaks indicate differential accessibility (Buenrostro et al.).

We performed ATAC-seq in control morphant and KMT5B morphant animal caps in triplicate across three biological replicates. We identified 118 809 total peaks, a result comparable to previous ATAC-seq experiments in Xenopus animal caps (Esmaeili et al.) (Figure 5A). In total, 24604 (20.7%) are found at promoters, 1648 (1.4%) immediately downstream of the TSS, 1512 (1.3%) at the 5’ UTRs, 996 (0.8%) at the 3’ UTRs, 4489 (3.8%) in the exons, 42188 (36%) in the introns, and 43372 (36%) in the intergenic regions. Of these peaks, 556 are differentially accessible (Figure 5B), representing 0.05% of all peaks. Surprisingly, most of the peaks were increasing in accessibility, while only 17 peaks became less accessible. Upon manual inspection of the local browser tracks, the less accessible peaks seemed to largely be called as the result of higher peaks in one of 3 control morphant injected tracks and may not have biological relevance. Contrary to our expectations, we did not find chromatin accessibility to change significantly on ciliogenic genes. When we look at 3 of the most misregulated genes from the KMT5B knockdown RNA-seq experiment, ttc23.L, dnal4.S, and ift88, their accessibility does not change (Figure 5C). The most significantly changing peaks are found mostly in intergenic regions (Figure 5D) and are generally increasing in accessibility rather than decreasing. Overall, this does not support the hypothesis that KMT5B exerts its control on ciliogenic gene expression through altering chromatin accessibility, and this regulation must occur through a different mechanism.

**Figure 5:**
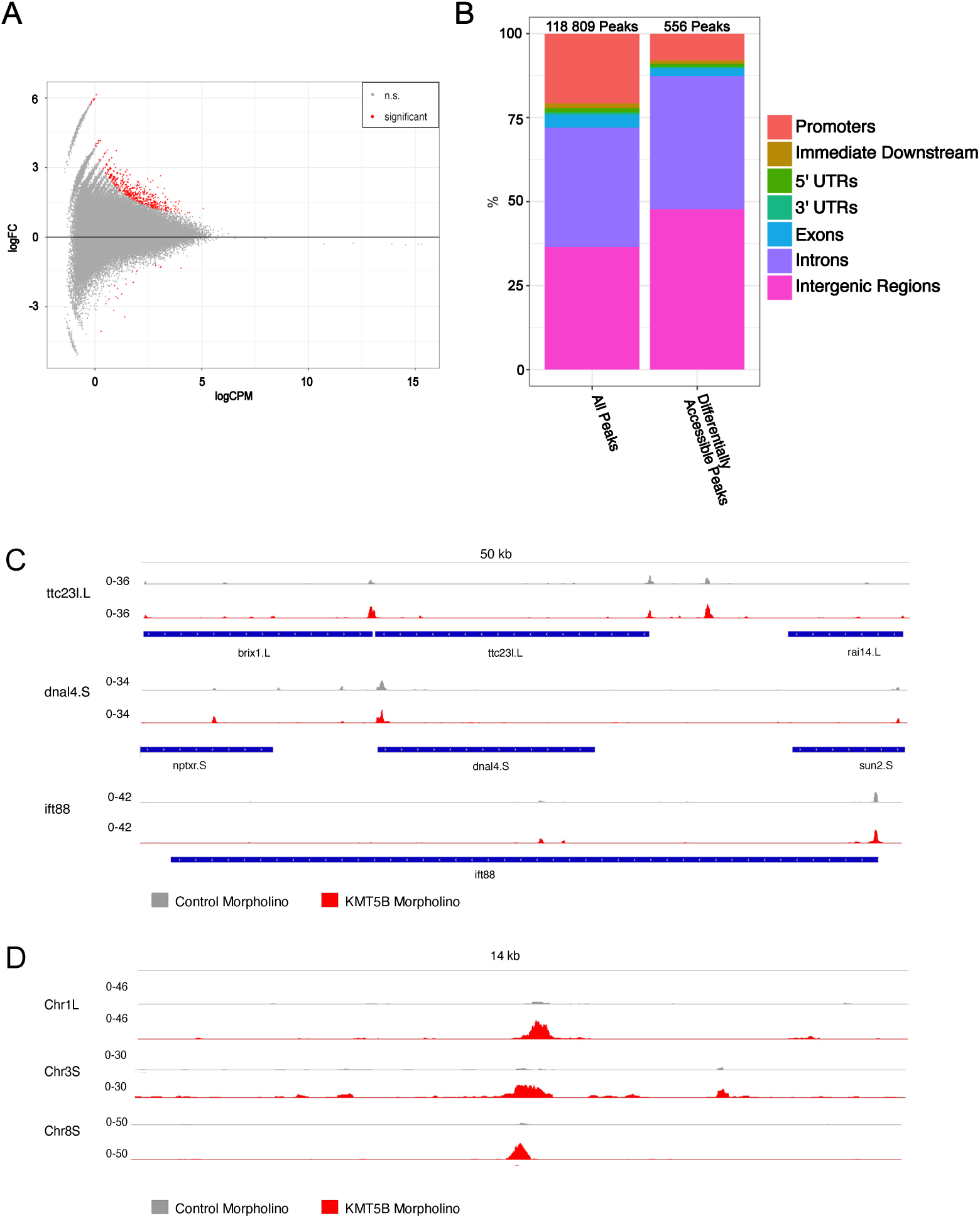
Depletion of KMT5B leads to increased chromatin accessibility. A) MA plot showing the distribution of peaks in KMT5B knockdown animal caps. Significantly changing peaks are shown in red (p<0.05). The majority of peaks are increasing rather than decreasing, indicating a mild chromatin opening effect of KMT5B knockdown in *Xenopus laevis* animal caps. B) Distribution of peaks across genomic features in all peaks and changing peaks. N = 3 biol. replicates. C) Representative browser tracks of the most significantly downregulated ciliogenic genes from the KMT5B knockdown RNA-seq experiment. D) Representative browser tracks of the most significantly changing peaks between the KMT5B knockdown and control animal caps.

## DISCUSSION

Multiciliated cells are a highly specialized and terminally differentiated cell type, that represent a critical component of mucociliary epithelia found on the human alveoli and the embryonic skin of *Xenopus* (Walentek; Werner and Mitchell; Jennifer L. Stubbs et al.). Multiciliogenesis is a complicated cellular differentiation process by which epidermal cells generate hundreds of motile cilia and this is executed through a complex gene expression program (Meunier and Azimzadeh; J. L. Stubbs et al.). Here we report that the H4K20 methyltransferase KMT5B regulates *Xenopus* multiciliogenesis. We show that KMT5B knockdown in animal caps leads to the concerted downregulation of ciliogenic genes, even when the master regulator of multiciliogenesis is overexpressed.

We have previously shown that the ciliogenic phenotype is specific and dependent on the catalytic activity of KMT5B. We generated the phenotype using two non-overlapping morpholinos and demonstrated that only catalytically active KMT5B mRNA is able to rescue (Angerilli et al.). Here we show that rescue with KMT5B demethylase, PHF8, also relies on catalytic activity. PHF8 is a Jumonji C Domain (JmjC) containing histone demethylase. It is target to promoters by binding of H3K4me3 through its PHD finger, and the catalytic activity is carried out by the JmjC domain. PHF8 knockdown also leads to a number of craniofacial and cytoskeletal defects. In PHF8 depleted Zebrafish embryos, the pharyngeal arches are reduced or absent, and brain development is impaired (Qi et al.). PHF8 is also involved in regulating cell cycle progression (Liu et al.). Here we confirm another potential function for PHF8 in multiciliogenesis and show that the rescue of cilia in KMT5B depleted organoids requires catalytic activity.

We report that two enzymes involved in H4K20 methylation, KMT5B and PHF8 have an effect on multiciliated cells that depends on catalytic activity. What remains unclear is whether they exert this function through methylation of H4K20. H4K20me1 has an ambiguous role in transcription. In some contexts, it seems to be a transcriptional activator (Shoaib et al.; Barski et al.), while in other contexts it is a transcriptional repressor (Congdon et al.; Gurvich et al.; Trojer et al.). The mark is largely found in the gene bodies of active genes. ChIP and ATAC-seq data also correlate H4K20me1 to the transcription of short housekeeping genes (Shoaib et al.). However, SET8 depletion results in a two-fold increase in the expression of H4K20me1-decorated genes (Congdon et al.). H4K20me1 reader, L3MBTL1 has also been linked to transcriptional repression, and demethylation of H4K20me1 by PHF8 derepresses a subset of E2F1-regulated promoters (Gurvich et al.; Trojer et al.; Boccuni et al.). Altogether, the role of H4K20me1 in transcription is unclear, and may be context- or location-specific.

Despite the fact that KMT5B and KMT5C both contribute to the writing of H4K20me2, only KMT5B has a function in ciliogenesis, and broadly in transcriptional regulation. If this function is exerted by controlling H4K20me1 levels, it may be that the effects of KMT5B and KMT5C are spatially segregated. KMT5C stably associates with heterochromatin, while KMT5B does not and presumably writes throughout the genome (Hahn et al.). While this would not stop KMT5B activity at heterochromatin, it may have higher activity in the more accessible euchromatic regions. This could explain the strikingly different effects of two similar enzymes on transcription and could be further explored in the future by mapping novel H4K20me1 peaks by CUT&RUN to determine whether a partitioning effect is involved in regulating ciliogenesis.

Alternatively, the ciliogenesis phenotype may not operate through altering the chromatin. Many histone modifying enzymes have non-catalytic functions, or may methylate non-histone substrates. Since catalytic activity of both KMT5B and PHF8 is required for regulating ciliogenesis, it could be that both of these enzymes are methylating non-histone targets. KMT5B has previously been shown to methylate in vitro non-histone proteins including CASZ and OSBPL1A (Weirich et al.), and SET8 has been shown to methylate α-tubulin (Chin et al.). Additionally, recent data from *Drosophila* with H4K20 histone mutants suggests that H4K20 writer and reader proteins SET8 and L(3)MBT may not require H4K20me to carry out their functions (Crain et al.). This could also be the case for other H4K20 modifying enzymes like KMT5B.

We previously demonstrated that KMT5B knockdown cannot be rescued by ovexpression of MCI (Angerilli et al.). In wildtype embryos, multicilin overexpression is sufficient to induce multiciliogenesis in non-MCCs. The promotion of MCC specification by multicillin is maintained in KMT5B knockdown embryos, based on the unambiguous presence of multiplied basal bodies, however these cells still lack cilia tufts. On the transcriptional level, overexpression of mci in control morphant animal caps led to a significant upregulation of ciliogenic genes, while these genes were significantly downregulated when multicilin was overexpressed in KMT5B knockdown caps. The changes in transcription are largely driven by KMT5B knockdown, not multicilin overexpression, despite the fact that multicilin overexpression leads to a dramatic remodelling of the embryo. Taken together this suggests that regulation of multiciliogenesis by KMT5B is achieved through an alternative pathway to the canonical multiciliogenic program.

But how is this regulation achieved? We initially hypothesized that aberrant hyperaccumulation of H4K20me1 on ciliogenic genes was the cause of this concerted downregulation. However, since mapping H4K20me1 by ChIP-seq was not technically feasible, we decided to pursue an alternative strategy. Because H4K20me1 has been linked to chromatin accessibility, we thought that KMT5B may control ciliogenesis by altering accessibility, either at ciliogenic genes, or as a whole. However, profiling accessibility by ATAC-seq showed only a mild effect on the chromatin. Only 0.05% of peaks were affected, and they tended to actually increase in accessibility rather than decreasing. Accessibility in the regulatory regions of ciliogenic genes was not affected. In fact, very few differentially accessible peaks were mapped to genes, and were instead found in intergenic and intronic regions. Overall, this suggests that KMT5B does not exert its regulatory role through alteration of chromatin accessibility (Shoaib et al.; Barski et al.).

We previously showed that KMT5B is responsible for writing approximately 50% of H4K20me2, with the remaining 50% being written by KMT5C (Angerilli, Tait et al. 2023). Following up on this finding, new data here indicate that mono- or dimethylation of H4K20 has little effect on chromatin accessibility in a global or significant manner. Regarding the function of KMT5B in ciliogenesis, it is still unclear whether this process is regulated through methylation of H4K20 or through an additional function of the enzyme itself. Our findings may suggest the presence of an unknown reader of H4K20me2 that drives ciliogenesis through deposition of the mark. Alternatively, KMT5B may carry out its role in ciliogenesis separate from H4K20 methylation, possibly through the methylation of a yet-unknown non-histone target. KMT5B is an essential gene, with knockout leading to perinatal lethality in mice (Schotta et al., 2008). Further research is required to establish the mechanism underlying these essential functions, including its role in ciliogenesis.

## MATERIALS AND METHODS

### Ethics statement

*Xenopus laevis* and *Xenopus tropicalis* were acquired from Nasco and Xenopus1. *Xenopus* experiments adhere to the Protocol on the Protection and Welfare of Animals and are approved by the local Animal Care Authorities (license number: 03-22-042).

### Xenopus methods

*Xenopus laevis* adults and embryos were handled using standard protocols as described in (Showell and Conlon). 2-cell stage embryos were injected with up to 5 nl per blastomere, and volume was scaled based on embryonic stage. 8-cell stage embryos were injected with 1.25 nl per blastomere. 2-cell stage embryos were co-injected with Alexa-488 Dextran (Invitrogen) and fluorescently stained embryos were co-injected with Hyls1-GFP, both as lineage tracers. 2-cell stage embryos were sorted left- and right-injected based on fluorescence. Embryos were cultured in 0.1x MMR at temperatures ranging from 16-23° and staged based on the Nieuwkoop Faber table of *Xenopus* development (NF 1967).

Animal caps were manually dissected by excising the blastocoel roof of blastula stage embryos (NF9). Animal caps were then transferred to a petri dish with individual wells to avoid amalgamation. The animal caps were incubated in Steinberg’s solution. MCI-hGR injected animal caps and embryos were induced at the midgastrula stage (N11) with 10 µM dexamethasone (Sigma).

### Expression constructs and morpholino oligonucleotides

Translation-blocking morpholino oligonucleotides directed against KMT5B (*X. laevis* KMT5B 5′-ggattcgcccaaccacttcatgcca-3’), KMT5C (*X. laevis* KMT5B: 5′-ttgccgtcaaccgatttgaacccat-3’) and standard control morpholino (5′-cctcttacctcagttacaatttata-3′) were obtained from Gene Tools LLC. In total, 30-40 ng of morpholino were injected per blastomere into 2-cell stage *X. laevis* embryos. For confocal analysis, embryos were injected at the 8-cell stage with 5 ng of morpholino into the ventroanimal blastomere.

Rescue experiments were performed with wildtype truncated human PHF8 (1-489) and catalytically inactive truncated human PHF8 (1-489) in pCS2+, kindly provided by R. Shiekhattar (Fortschegger et al.). MCI-hGR in pCS2+, kindly provided by P. Walentek. Further, we injected Hyls1-GFP in pCS2+ kindly provided by A. Dammerman, and LacZ in pCS2+ as lineage tracers. Lastly, we injected luciferase mRNA as a control. Synthetic mRNAs were injected at the 2-cell and 8-cell stage.

### Immunocytochemistry

Wholemount immunocytochemistry was performed as previously described (Robinson and Guille) on tailbud stage embryos (NF28). Confocal stainings were performed the same way, with the exception of skipping the methanol step and permeabilizing the cell membranes with a 20-minute incubation in PBS with 2% Triton. Embryos were incubated with monoclonal anti-acetylated α-tubulin antibody (Signma-Aldrich, T6793 1:500) as a primary antibody, and an alkaline phosphatase-conjugated secondary antibody (sheep x mouse Fab IgG Alk Phos, Chemicon, 1:1000). For confocal staining, a fluorescent secondary antibody (goat anti-mouse 1:1000, Chemicon). Cell borders and the apical actin meshwork are stained by incubating in DAPI (1:50 in PBS, Sigma) for 15 minutes and in 5% Alexa Fluor 555 Phalloidin (Cell Signalling) in PBS for 1.5 hours. Embryos were mounted between two glass coverslips separated by 0.35 mm thick double-sided with wells cut into the tape and filled with modified Dabco mounting solution (85.43% Glycerol, 10% 10x PBS + 2% Dabco, 4.57% H_2_O). Samples were visualized on an inverted confocal microscope (Leica TCS SP8X).

### Statistical analysis

ICC results from knockdown experiments were analysed using a two-tailed Student’s t-test. Results from rescue experiments were analysed using one-way ANOVA with post-hoc Tukey test. Asterisks indicate p-values: * = p < 0.05, ** = p < 0.01, *** p < 0.001.

### RNA library preparation and sequencing

Ten *Xenopus laevis* animal caps per condition were harvested and snap frozen on liquid nitrogen. RNA was extracted using the RNeasy Mini Kit with on column DNAse digestion (Qiagen). Total RNA was measured on a 4200 Tapestation (Agilent) and the quality was checked to ensure a RIN^e^ of at least 7. Library preparation was performed using NEBNext®UltraTM II Directional RNA Library Prep Kit for Illumina®(New England Biolabs). PolyA(+) mRNA was selected for using the NEBNext®Poly(A) Magnetic Isolation Module (New England Biolabs). NEBNext®Multiplex Oligos for Illumina®(Index Primers Set 1 and 2) were used for sample indexing. Size selection was performed using AMPure XP beads (Beckman Coulter). After preparation, the finished libraries were rerun on a 4200 Tapestation (Agilent) using the HSD1000 ScreenTape (Agilent). Sequencing was performed on an Illumina NextSeq1000 with 50bp paired-end reads to a depth of 20 million reads.

### RNA-seq analysis

All data processing methods were applied using default parameters unless specified. Expression quantification was performed using kallisto (version 0.48) using XENLA_10.1 version of the Xenopus laevis genome. In R/Bioconductor, expression data were collapsed from isoform to gene level for downstream processing. Differential expression was assessed using DESeq2 (version 1.42.1) using the experimental batch as random factor. RNA high-throughput sequencing data has been deposited in the NCBI GEO under the accession numbers GSE274392.

### ATAC library preparation and sequencing

Two animal caps per condition were cultured until neurula stage (NF16) and collected for ATAC seq. Samples were prepared as in (Buenrostro et al.; Esmaeili et al.). In short, embryos were centrifuged for 5 minutes (500 g, 4°C) in PBS. The PBS was replaced with lysis buffer (10 mM Tris-HCl (pH 7.5), 4 mM MgCl2, 10 mM NaCl, 0.1% Igepal CA-630) and animal caps were homogenized by pipetting. The tagmentation reaction was perfomed in 25 µl TDE1 buffer, 1.88µl TDE1 enzyme (Illumina 20034197) diluted to a final volume of 50 µl with nuclease-free H_2_O at 37°C with 350 rpm shaking for 30 minutes. Libraries were prepared according to Buenrostro et al. and assessed using a Tapestation (Agilent) before and after bead-based size selection. Samples were sequenced with 50 bp, paired end reads to a depth of 20 million reads per sample on an Illumina NextSeq 1000.

### ATAC-seq analysis

All data processing methods were applied using default parameters unless specified. After adapter trimming with cutadapt, sequencing reads were aligned to the XENLA_10.1 version of the Xenopus laevis genome using bowtie2 (version 2.4). Duplicates were marked and removed with Picard Tools. Peaks were called using macs2. Aligned reads were read in R/bioconductor, converted to coverages (library GenomicRanges), and exported to bigWig files (library rtracklayer). Differential accessibility was assessed using csw/edgeR and TMM normalization as described in the csaw workflow detailed in https://github.com/reskejak/ATACseq/blob/master/csaw_workflow.R. ATAC-seq data as been deposited in the NCBI GEO under the accession number GSE274391.

## ACKNOWLEDGEMENTS

We thank Drs. Peter Walentek, Alexander Dammerman, and Ramin Shiekhattar for their kind gift of recombinant plasmids; Dr. Andreas Thomae and the Core Facility of Bioimaging of the Biomedical Center, LMU Munich for the instructions and technical support in confocal imaging, and the BMC Core Facility Animal Models (CAM) for animal care and maintenance. This work was funded by the Deutsche Forschungsgemeinschaft (DFG, German Research Foundation) – Project-ID 213249687 – SFB 1064 (Project A12).

## Supplementary Figures

**Figure S1:**
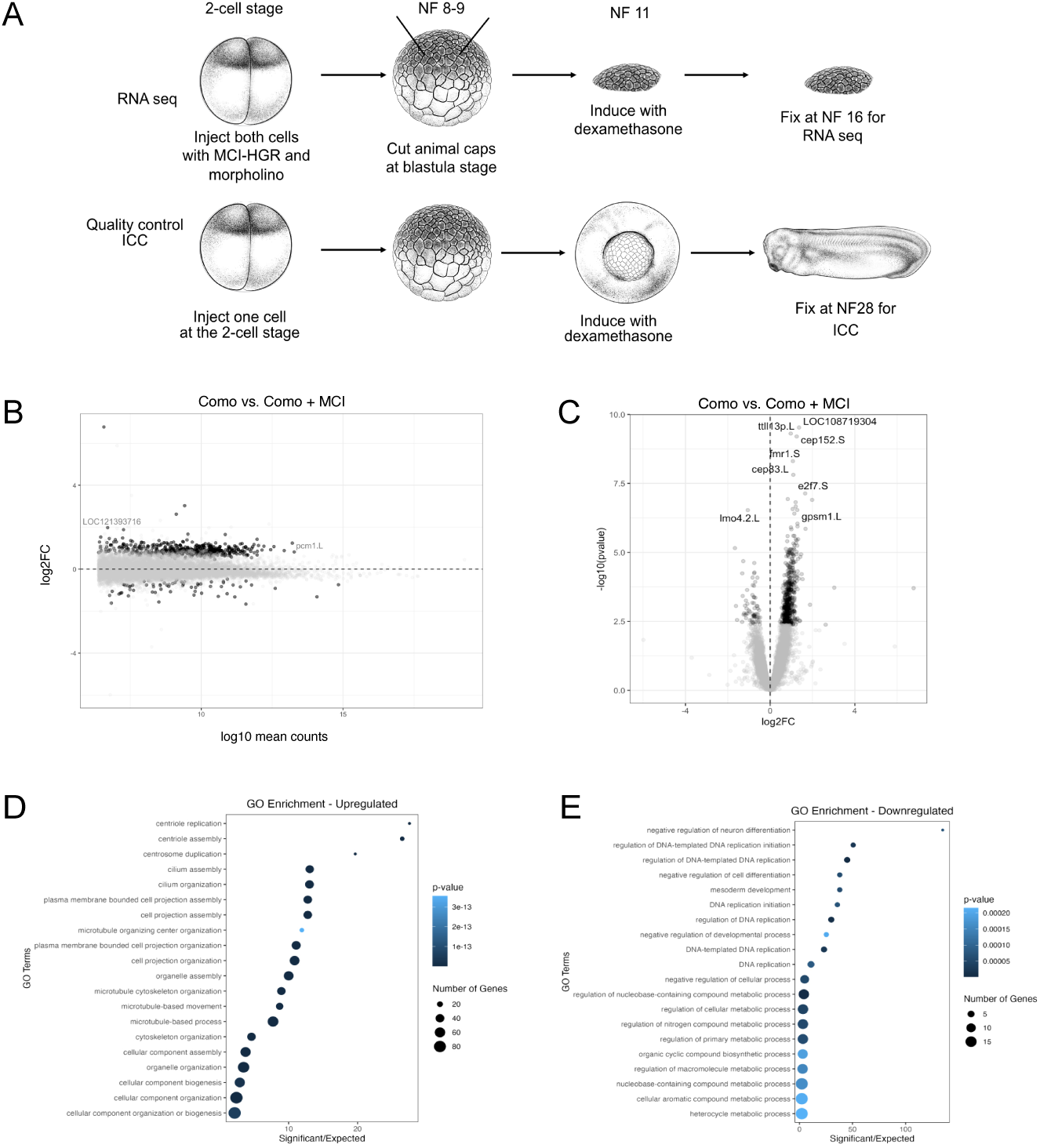
Overexpression of MCI leads to upregulation of multiciliogenic genes. A) Protocol for multicilin overexpression. Embryos are injected in both cells of the 2-cell stage with the morpholino and mRNA of interest. They are allowed to develop until the blastula stage (NF8-9), at which point animal cap explants are dissected. At late blastula stage (NF 11), animal caps are induced with 10 µM dexamethasone. Animal cap explants are harvested at the neurula stage (NF16) and further processed for RNA-seq. Simultaneously, sibling embryos are injected on one cell of the 2-cell stage with constructs of interest and allowed to develop until late blastula stage (NF11). Embryos are then induced with 10 µM dexamethasone. At tailbud stage (NF28), embryos are fixed for whole mount immunocytochemistry. These quality control embryos are assessed to exhibit both a KMT5B knockdown phenotype and a multicilin overexpression phenotype. B, C) MA plot (B) and volcano plot (C) showing the change in gene expression in control morpholino + MCI-hGR overexpressing animal caps. D, E) GO analysis showing the upregulated (D) and downregulated (E) gene categories upon MCI-hGR overexpression. The upregulated gene categories are largely related to ciliogenesis and microtubule assembly.

